# Hook shape of growing leaves results from an active regulation

**DOI:** 10.1101/2020.02.27.967513

**Authors:** Mathieu Rivière, Yoann Corre, Alexis Peaucelle, Julien Derr, Stéphane Douady

## Abstract

The rachis of most growing compound leaves observed in nature exhibit a stereotyped hook shape. In this study, we focus on the canonical case of *Averrhoa carambola*. Combining kinematics and mechanical investigation, we characterize this hook shape and shed light on its establishment and maintenance. We show quantitatively that the hook shape is a conserved bent zone propagating at constant velocity and constant distance from the apex throughout development. A simple mechanical test first reveals non-zero spontaneous curvature profiles for the growing leaves, indicating that the hook shape is actively regulated. It then evidences the robust spatial organization of growth, curvature, rigidity and lignification and their interplay. Regulation processes appear to be specifically localized: in particular, differential growth occurs where the elongation rate drops. Finally, impairing the graviception of the leaf on a clinostat led to reduced hook curvatures but not to its loss. Altogether our results suggest a role for proprioception in the regulation of the apical hook, likely mediated via mechanical strain.

## Introduction

Plants display an impressive variety of shapes. This diversity can be observed across different species or organs, but not only. Single organs also adopt different shapes during their development. In this wide diversity of shapes, some stand out by their uniqueness or their beauty while some others stand out by their ubiquity. Ubiquitous shapes are of particular interest, overall when observed at developmental stages. Since there exists a strong and intimate link between the shape of a developing organ and its growth kinematics, such ubiquitous shapes could suggest shared processes of shape and growth regulation among species.

The apical hook is a perfect example of such shapes. It consists in a marked downward bending of growing organs near their apical end. The term ‘apical hook’ is usually associated with the hypocotyls of dicots. However many stems–with opposite phyllotaxis–and pinnate or bipinnate leaves display a similar shape (see Fig. 1). The growing fronds of some ferns, Cycas or Droseras, exhibit an even more exaggerated curvature at the apical end of their fronds, referred to as circinate vernation. In each case, the apical hook is associated with a characteristic opening, unrolling or unfurling motion that keeps going until the organ reaches its mature state (1, 2).

**Fig. 1.**
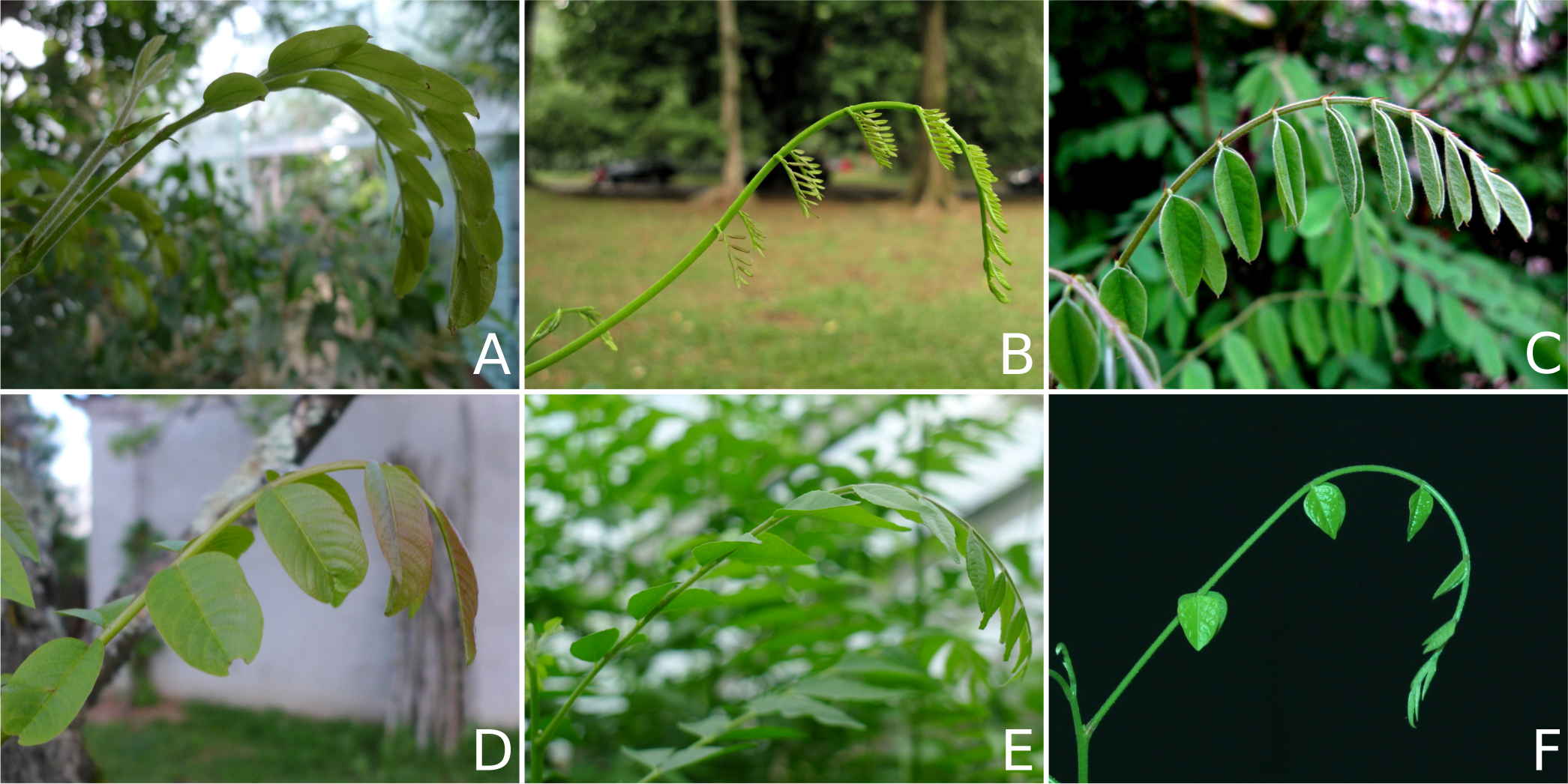
The typical hook shape of the rachis found in growing compound leaves illustrated in six different genera. (A) *Calliandra haematocephala*, (B) *Sindora wallichii*, (C) *Amorpha fruticosa*, (D) *Juglans regia*, (E) *Murraya koenigii*, (F) *Averrhoa carambola*, our plant model in this study. Images (B) and (C) taken from the supplemental information of (1) where more examples can be found.

The apical hook has mostly been studied in hypocotyls since the founding work of Charles and Francis Darwin (3). It has since remained ‘one of the favorite experimental models to study the regulation of differential growth’ and is up to this date an active domain of research (4). In this study we will rather focus on the case of growing compound leaves, in which–as far as we know–the hook shape remains an open problem. More precisely, we will here work on *Averrhoa carambola* which leaves are odd-pinnate and display a marked apical hook during their development.

This study originates in a naive observation: it looks like growing compound leaves droop under their own weight. Is this true, though? The shape of the leaf could either be the result of a passive mechanical buckling or of an active posture regulation process. In the latter case, the question of the regulated quantity immediately arises. Is the curvature of the hook regulated, or rather the spontaneous curvature that it would have without gravity?

Moulia et al. (5) have shown that the self-weight of mature maize leaves had a non-negligible impact on their shape. Could this also be valid for developing leaves? In such a case, the mechanics of the organ would dictate the final shape. These mechanical properties indeed change during growth. For instance, it is well-known that, like stems, compound leaves undergo a strong rigidification in the course of their development through lignification (6).

Plants are also sensitive to their environment. They can actively trigger growth-driven motions–called tropisms–to adapt their posture, following given external cues. Gravity and light are two obvious external cues, regularly put forward to explain plant shaping (7). However Bastien et al. (8, 9) have highlighted the importance of proprioception and autotropism in posture regulation processes. The latter works have recently inspired a model for the development of stem-like organs which involves gravitropism, autotropism and elasticity (10). From this model, many different realistic shapes could be predicted, suggesting that the combination of these three parameters packed the essential of posture regulation processes.

Here, the hook shape naturally questions the role of graviception and self-weight. In this paper we investigate experimentally the nature of the hook, passive and active, by combining a kinematics study with a simple yet rich mechanical test: the “flipping test”.

## Material and methods

### Culture conditions for *Averrhoa carambola* plants

Carambola seeds obtained from commercially available star fruits were grown in a small lab greenhouse. When sufficiently developed and robust, they were moved inside the experimentation room. There, plants have been submitted to a 12/12 light cycle under regular culture lighting lamps (ORTICA 200W 2700K). The temperature and relative humidity rate were measured. It was checked that no brutal variation of these quantities occurred during the experiments. The usual temperature values were comprised between 20 and 24 degrees celsius. The usual values for relative humidity were around 60%.

### Geometrical framework

Developing leaves were observed in the plane of the hook, that is to say thanks to side view takes. Rachis were reduced to their midlines and characterized within the geometrical framework depicted in Fig. 2. We define a global right-handed Cartesian coordinate system 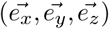 where 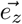 indicates the downward vertical. The plane of interest is defined by 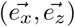 and we will refer to it as the principal plane. We define 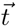, a local tangent vector to the rachis. The local orientation *θ* of the rachis is then given by 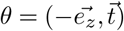. Finally we label the curvature of the rachis in the plane of interest *K*_*//*_. The rachis is furthermore described by its arc length s, from base to apex (or by the reverse arc length *s*_*R*_ from apex to base).

**Fig. 2.**
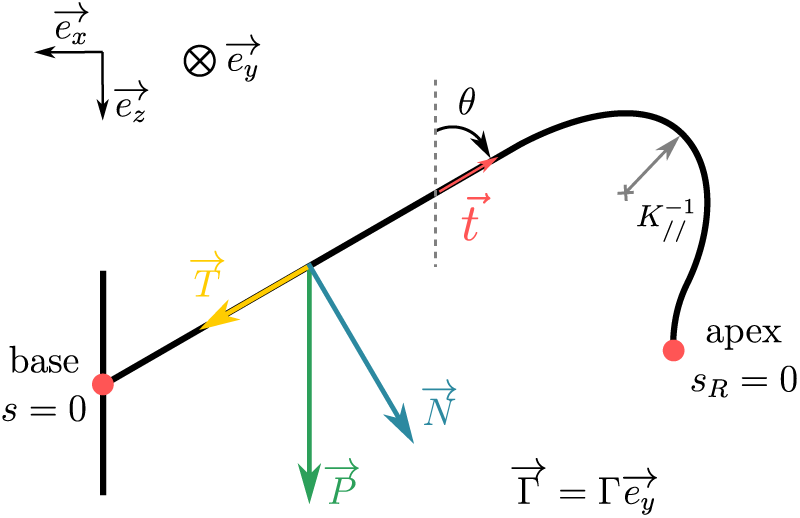
Geometrical and mechanical quantities describing our problem. We set our parametrization in the plane 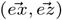 parallel to the hook (side view). The rachis is described by its arc length s (from base to apex) or equivalently by its reverse arc length *s*_*R*_ (from apex to base). The local orientation of the rachis is given by 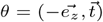 where is the vector tangent to the rachis. The local curvature is given by *K*_*//*_. Consider the weight applied at any point of the rachis. It is always possible to decompose it into the sum of two forces and respectively tangent and normal to the rachis. The sign convention for an arbitrary torque Γ is indicated.

### Geometrical characterisation of the rachis

The midline of the rachis has been extracted thanks to an algorithm based on Voronoi diagrams previously developed in the team (see (11) for more details). This technique has already proven efficient on cylindrical plant organs and is available as a MATLAB toolbox (12). In the case of compound leaves, however, leaflets make using Voronoi-based algorithms harder. Because they conceal a significant portion of the rachis and are of the same color, they considerably impact the shape of the retrieved skeleton. Depending on the experiment, additional procedures have been necessary to retrieve the shape of the rachis.

In experiments where the number of involved pictures is low (see later “flipping test”), the leaflets have been manually removed from the pictures with an image edition software. It was then possible to use our algorithm on the edited images. In other experiments where the number of pictures was too high (see later “growth kinematics”), the midline processing was split in two steps. First, an erroneous midline has been extracted using our Voronoi-based algorithm. Second, the midline was fitted with a set of Bézier curves. This procedure allowed to get rid of the skeleton errors due to the leaflets (see Fig. S1).

### Growth kinematics

We have accessed the growth kinematics with time lapse photography. The plant was set so that the principal plane (ex,ez) would be orthogonal to the optical axis of the camera (CANON EOs 1000D). A snapshot has been taken every 10 minutes. The elongation of the leaflet has been obtained by tracking the position of the leaflets. The evolution of the relative position of two successive leaflets indeed provides a coarse quantification of the strain rate (or relative elongation rate) of the rachis. The trajectories of the leaflets have been approximated to get a smoother estimation of the strain rate (see Fig. S2). At last, the complete strain rate field can be estimated by linear interpolation along the rachis, between leaflets (see Fig. S3).

### Mechanical quantifications and measurements

In some previous studies interested in quantifying the mechanical properties of plant organs, authors would actively bend the organs of interest (5, 13, 14) using loads. Here we do not affect the load but rather flip the arrow of self-weight itself. To extract the mechanical properties of the rachis we then resort to beam theory (13). A number of approximations are required, though. First, the section of the rachis is assumed to be circular. We also assume homogeneous density and isotropic mechanical properties. A reasonable approximation is *ρ* = 1 *kg* · *m*^−3^, typical of most biological soft tissues. At early stages of hook formation, we quantified the relative contribution of the leaflets to be less than 15% of the total weight. Because leaflets contributing to the torque (the one closer to the apex) are smaller, their contribution to the total torque is even less than 15%. Therefore, we neglect the mechanical action of the leaflets in this study. A side experiment performed after leaflets ablation confirms qualitatively that the presence of leaflets do not change the hook shape (see Fig. S4). Finally, the values of R were quite sensitive to the presence of trichomes. We have noticed that for each time step of our experiment the spatial profile of radii was approximately linear and steady. We thus performed a linear regression over all the collected values of R and used the fitted values in subsequent calculations.

Beam theory allows to link the torque–or moment of force–Γ applied on a beam to its flexural rigidity *B*, its curvature *K*_*//*_ and its spontaneous curvature *K*_0_ along the axis as follows (15):

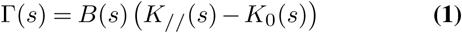

where *s* is the rachis arc length. The flexural rigidity *B* is furthermore related to the Young’s modulus *E* of the considered material by *B* = *EI* where *I* the second moment of area of the considered object. The flexural rigidity *B* and the spontaneous curvature *K*_0_ being intrinsic properties of the rachis, we see from equation 1 that we simply need two conformations of the rachis to evaluate them. This is made possible by the “up” and “down” orientations of the leaf during an experiment.

In our case, the self-weight of the rachis is the only mechanical action, the weight of the leaflets being neglected. Its intensity in any point of the rachis is estimated by measuring the radius *R* of the rachis and with the cylindrical approximation of the rachis’ geometry. It can then be shown that the intensity of the corresponding torque is obtained by spatially integrating the normal component of the self-weight.

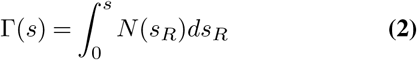

### A simple mechanical test: “the flipping test” - Protocol

A plant with a growing leaf is firmly fixed on a vertical clinostat^1^. In our case the clinostat was used to safely and consistently flip the plant upside-down in a dozen of seconds (i.e. a rotation of 180°). The other leaves are gently tied to the stem to prevent them from hiding the leaf of interest when flipped. Leaves have been maintained upside-down during 30 min and then released to ordinary orientation. This simple procedure has been repeated throughout the development of the leaf.

It is true that a gravitropic stimulus of a few minutes is sufficient to trigger gravitropism and posture correction in plants (see (16) for instance). However, the gravitropic response of a plant is proportional to the sine of the presentation angle (17, 18). In other words, no gravitropic response to the flip is expected. By comparison with undisturbed plants, we checked the development was not severely impacted. Some slow viscous relaxation of the plant appears after each conformation change. In the present study, we consider it as negligible (19). After having retrieved Γ and *K*_*//*_ for each of the two conformations (normal and flipped) through our image analysis, we can access the spatial profiles of the intrinsic mechanical parameters defined earlier, *B* and *K*_0_. At each point of the rachis, we have:

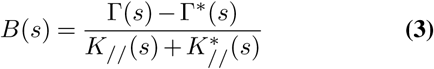

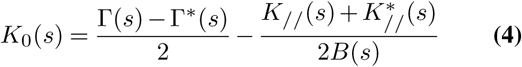

where the starred quantities refer to the flipped conformation of the rachis. At last, the repetition of this procedure for several days on the same leaf allows to monitor the evolution of these mechanical profiles throughout its development.

## Results

### Steady propagation of a bent zone

To put in context the unfurling motion of the leaf, we provide a 16 days time-lapse video (supplementary video 1). The first few days (until day 9), the new leaf slowly emerges, and eventually reverses to establish a hook shape (19). From day 10 onwards, the hook shape propagates from the base of the leaf to the apex while the leaf expands: it is the unfurling motion.

We quantify the unfurling motion by tracking the curvature both in space (along the rachis) and in time, and we represent it in a spatio-temporal diagram (see Fig. 3). We smooth the profiles *K*_*//*_ using a moving average of 5 h. Fig. 3 evidences two successive phases in the motion. From *t* = 0 *h* to 70 *h*, the rachis as a whole is curved. Curvatures decreases along the rachis to end with negative values at the apex. This first phase actually corresponds to the end of the hook shape establishment.

**Fig. 3.**
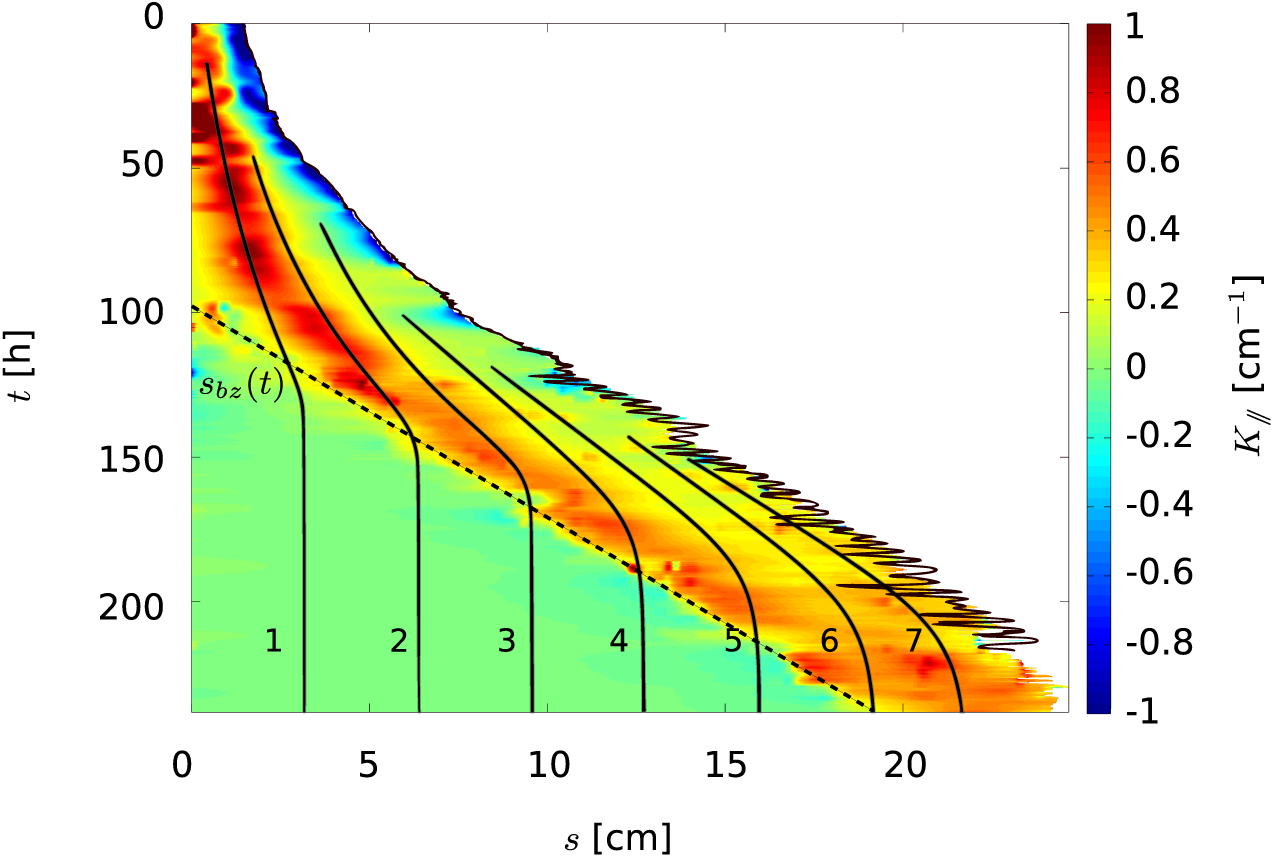
A typical spatio-temporal diagram of the rachis curvature *K*_*//*_. Note here that we use the regular arc length s to emphasize the growth of the leaf. On the right side of the diagram, the plain black line indicates the total observed length *L*(*t*) of the rachis. It shows some degree of oscillation corresponding to lateral movement of the leaf. The dotted line *s*_*bz*_(*t*) represents the delimitation of a bent zone: the hook. The translation of this bent zone corresponds to the unfurling motions described earlier. The lines numbered 1-7 correspond to the positions in time of the successive leaflets 1 to 7. Acquisition lasted for 250 hours (≃10.5 days). For the purpose of this graph, *K*_*//*_ profiles have been smoothed using a moving average with a 5h-window.

Then, from *t* = 100 *h*, we see the signature of the straightening, unfurling motion with unending of the basal end of the hook. It is now possible to distinguish two zones on the rachis: a bent zone and a straight zone. The straight zone expand while the apical end of the hook remains bent. This is why this process is often referred to as the ‘hook maintenance’ in the literature (4). This hook maintenance results in the translation of the hook along the rachis (often compared to a “fountain” (20)). From *t* = 110*h*, the position of the bent zone with respect to the apex is almost constant. Its position *s*_*bz*_(*t*) is linearly fit with a dotted line in Fig. 3 with slope *v* ≃ 1.3 *mm* · *h*^−1^. This unfurling motion lasts until the developing leaf reaches complete straightness. Note that we considered here that the motion is purely two-dimensional^2^.

### Colocalization of growth and bending

The trajectory of the leaflets are well fitted by a logistic-like curve (see Fig. S2) giving a good indication of the growth field. For each of the leaflets, its displacement is always taking place in the previously defined bent zone (see Fig. 3). For instance, the interleaflet defined by trajectories 1 and 2 goes through an elongation phase between *t* = 0*h* and *t* ∼ 120*h* before reaching its final length. Then, we see that the closer to the apex, the later a given interleaflet begins its elongation phase and reaches its final length. The most striking observation of this colocalization is the fact that the leaflets trajectories becomes completely vertical (meaning no growth) once they arrive in the zone *s* < *s*_*bz*_ (no curvature). Conversely, in the curved region *s* > *s*_*bz*_, the leaflets trajectories are evolving in time, corresponding to a growth zone (see Fig. S3). These results show that the bending zone and the growth zone overlay.

### Self-weight contribution to shape is negligible at earliest stages

We present in Fig. 4 the results of the ‘flipping’ experiment obtained on a single young leaf. As described in the Materials Methods section, the leaflet is flipped upside-down in a few dozens of seconds. Pictures are taken before the flipping (see Fig. 4A) and as soon as the 180° rotation is over (see Fig. 4B). For the sake of comparison, Fig. 4B has been rotated and overlaid with Fig. 4A by maximum intensity projection (see Fig. 4C). Arrows indicating the position of the apex in normal (blue) and flipped (yellow) configurations have been added to make the comparison easier. As it appears in Fig. 4C, the shape of the rachis barely changes when tilted. As expected, the rachis as a whole is slightly displaced upwards. This displacement is small though: it is negligible in comparison with the total length of the rachis, and smaller than its radius. The same flip experiment was repeated for a week on the same leaf (see Fig. 5A). It shows that the deformation of the rachis due to the flip increases during the course of development, as the length and weight of the leaf. To quantify this effect the spontaneous curvature *K*_0_ and the flexural rigidity *B* were extracted for each repetition of the flip experiment (see Materials Methods), and the results are shown in Fig. 5B and Fig. 5C respectively.

**Fig. 4.**
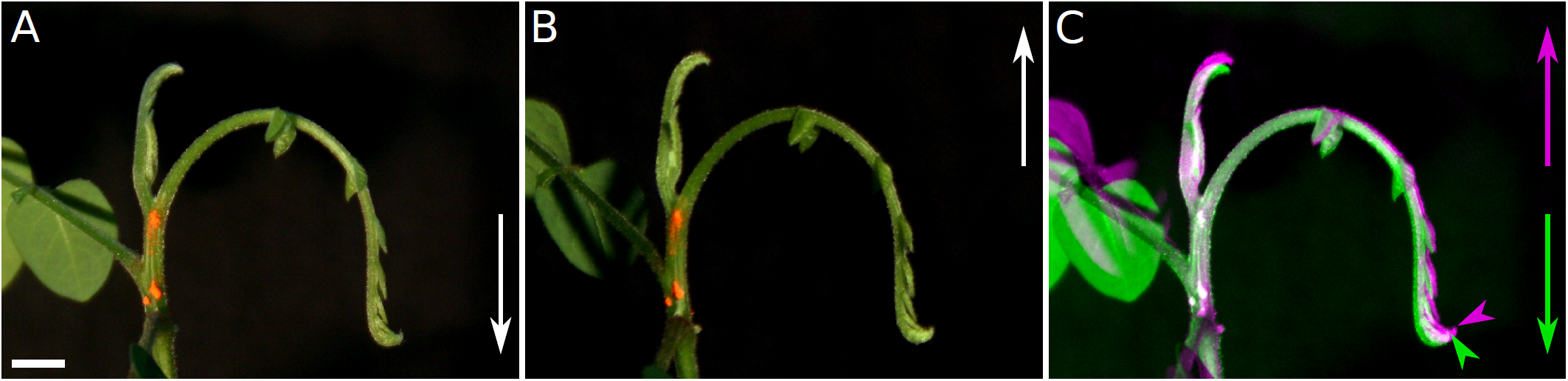
Self-weight of the leaf has a negligible impact on its shape. (A) The young leaf in the regular orientation. The acceleration of gravity g is indicated. (B) The same leaf flipped upside-down. (C) False color overlay of (A) and (B) flipped. Green color corresponds to (A), purple to (B). Scale bar = 1 cm

**Fig. 5.**
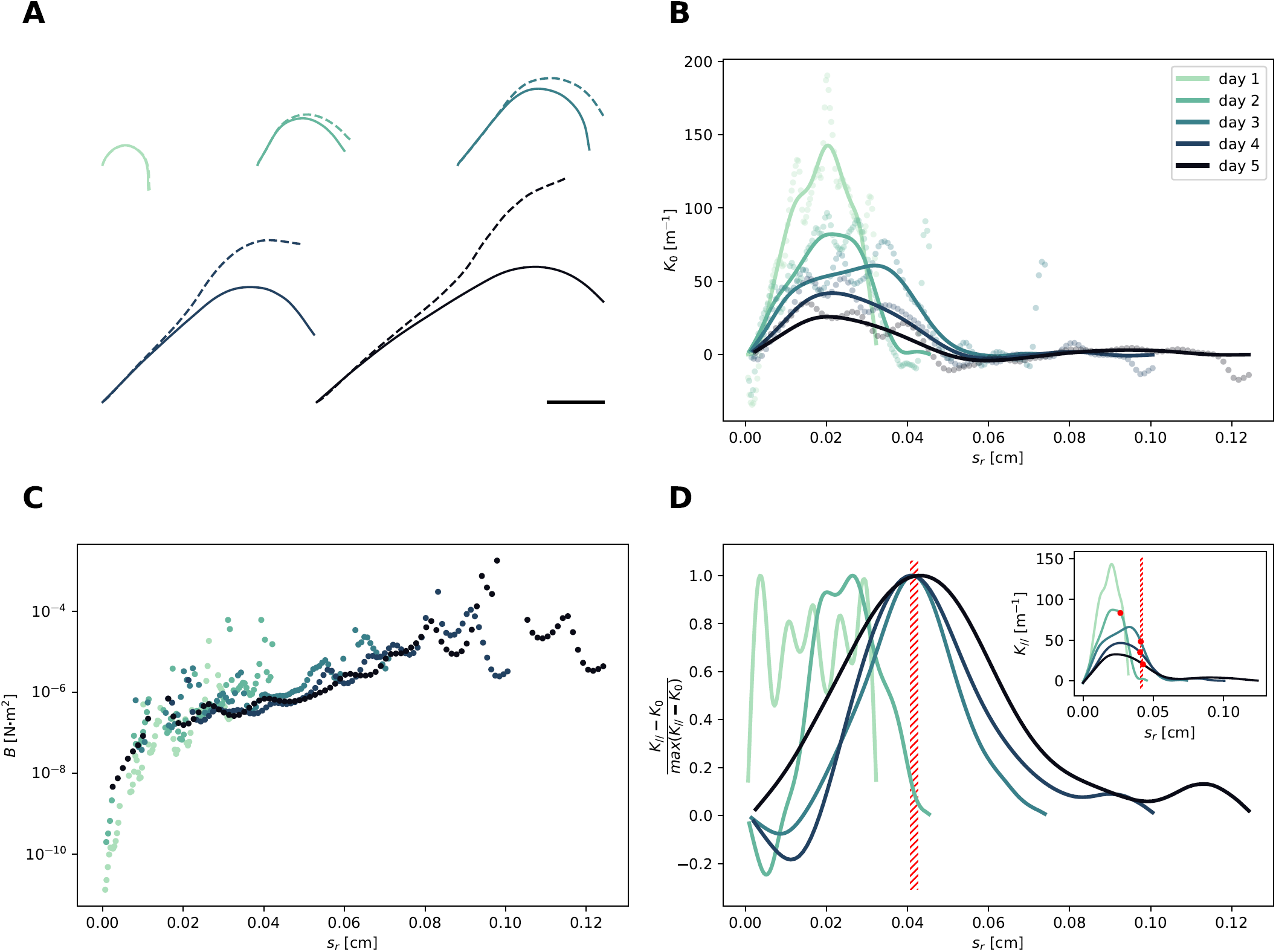
Mechanical properties of the rachis during development. (A) For each day of the experiment (left to right), the skeletons corresponding to the regular (plain line) and flipped (dotted line) orientations are overlaid. Scale bar = 2 cm. (B) Spontaneous curvature *K*_0_ along the rachis. Dots and lines respectively correspond to a moving average of size 1% and 6% of the leaf length. (C) Flexural rigidity *B* along the rachis. The arclength used here is the reverse arc length *s*_*R*_ (i.e. the distance to the apex). Data shown here were obtained from *K*_0_ values with a 1% smooth. (D) *K*_*//*_ − *K*_0_ normalized by its maximum. The red hatched zone shows the position of the maxima for the three last days of experiment. Inset: *K*_*//*_ along the rachis. Red dots indicate the exact positions of the maxima.

### Mechanical test reveals spontaneous curvature profiles for the growing leaves

Fig. 5B evidences spontaneous curvature near the apex. It also shows that the spontaneous curvature maintains throughout growth, even if with a decreasing intensity and increasing extension toward the base. At first, the spontaneous curvature is large *K*_0_ = 240 *m*^−1^ but drops to 140 *m*^−1^ in one day and keeps on decaying, but it is somewhat compensated by the increase in the width of the curved region. As a result the hook gets larger while keeping a nearly constant vertical downward angle at the apex.

### Flexural rigidity profile appears to be steady in time

Fig. 5C shows the spatial profiles of the flexural rigidity *B* for each day of the experiment. We see that *B* spans on a wide interval. The vast majority of values lies between nearly 10^−7^ *N* · *m*^2^ and few 10^−5^ *N* · *m*^2^, that is to say 2 to 3 orders of magnitude. There is a visible tendency in the profile of *B*: the flexural rigidity strongly increases from apex to base. In contrast with the case of *K*_0_, we notice that all the *B* profiles are collapsing on a single ‘master curve’. In other words, the profile of *B* does not vary much on the duration of the experiment.

### Deformation due to gravity is maximal downstream of bending zone

From the previous quantities we draw *K*_*//*_ − *K*_0_, the contribution of gravity to curvature in regular conditions. Fig. 5D shows *K*_*//*_ − *K*_0_ normalized by its maximum. We observe that after a shift at the beginning, this profile also tends toward a constant shape and localization. Note that *K*_*//*_ − *K*_0_ is proportional to the strain rate of the rachis under its own weight. Remarkably, the maximum of *K*_*//*_ − *K*_0_ is set in a zone of decreasing *K*_*//*_, even when a steady profile is not reached (see red hatchered zone, inset).

### Hook shape persists even with di*s*_*R*_upted graviception

By using a clinostat, we compared the development of leaves grown with impaired graviception to leaves growing in a constant gravity field. The spatio-temporal graphs of *K*_*//*_ for both leaves are shown in Fig. 6. For a leaf grown in regular conditions (see Fig. 6A), we recover the typical behaviour of the unfurling motion defined earlier. In particular, we can see a well defined bending zone near the apex. Similarly, for a leaf grown on the clinostat (see Fig. 6B), we see that a bending zone also exists. Its propagation is qualitatively comparable to the propagation in the regular case. Down-stream of the bending zone, the rachis tends to become flatter and flatter, just as in the regular case. Though, the mean curvature of the bending zone is significantly lower in the case of the leaf grown on the clinostat (typically 2 to 3 fold less than for the regular case).

**Fig. 6.**
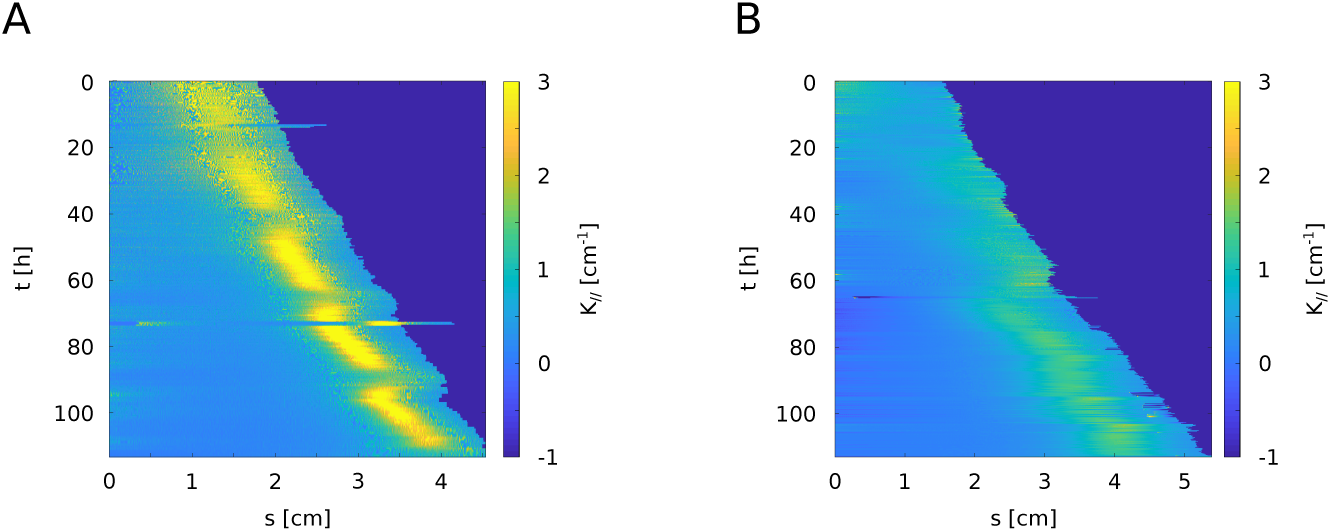
Influence of graviception on the hook. Spatio-temporal evolution *K*_*//*_ of for two different leaves with the same plastochrone index. (A) Leaf grown in regular conditions. This leaf displays a bending zone characteristic of the unfurling motion. (B) Leaf grown on the clinostat. Even with impaired graviception, a hook is visible, with lower curvature though.

## Discussion

### The unfurling motion and the related hook are actively regulated

A naive look at the hook would lead to think that the leaf droops under its own weight, somewhat in the fashion of mature maize leaves (5). We have demonstrated with a simple experiment that this is not the case. During the first developmental stages of the leaf, the elastic deformations of the rachis due to the self-weight of the leaf do not contribute to its shape (see Fig. 4). The spontaneous curvature of the rachis is non zero (see Fig. 5B). This confirms the active character of the hook shape. In other words, compound leaves do not droop but actively grow curved, at least during the first stages of the development of the leaf. This result is not trivial if one considers the leaf from a static point of view. It makes more sense once one considers the dynamic aspect of the leaf unfolding mechanisms where the curvature is evidenced to evolve continuously, even changing sign, to finally achieve a flat state.

### Regulation processes appear to be precisely localized

The unfurling motion occurring during the development of *Averrhoa carambola* pinnate leaves lies on specific kinematics (see Fig. 3). A steady bending zone with a well-defined length runs from the apex. It propagates at approximately a constant velocity. This kind of “fountain” kinematics recalls the maintenance of the apical hook in the case of hypocotyls (2, 20). While the active character of the uprising of the rachis–that is the unfurling motion strictly speaking–is not controversial for the least, the question remained open in the case of the hook. Instead, we have shown that the robust maintenance of the hook lies in the fine regulation of several quantities.

First, we have evidenced that the rachis grows mostly in the bent zone, near its apex (see Fig. 3). The hook is thus colocalized with active processes in the first place. The elongation and bending zones largely overlap, implying that the hook is a “steady from [obtained] from changing cells” (21).

Second, we have put forward a steady profile of flexural rigidity, with a gradient from apex to base (see Fig. 5C). Although the measurements of *B* are very sensitive to curvature measurements (and thus noisy), they are consistent with rigidity gradients observed in mature maize leaves (5). This flexural rigidity gradient is partly consistent with the increase of radius along the rachis. Geometry contributes to the variations of *B*, but because *B* scales like *R*^4^, and the radius *R* was found to vary by about a factor 1.25 the geometrical variation is only about 1.25^4^ = 2.5. The other factor is the gradient of lignification: we performed complementary experiments to qualitatively reveal this lignification gradient (see Fig. S5). Our results are consistent with previous findings showing an increase in the Young modulus of cell walls by two to three orders of magnitude (6).

It appears that the uprising motion occurs in a region of the rachis which is the seat of numerous regulations: flexural rigidity, spontaneous curvature, end of growth, beginning of lignification. This region also happens to be the region where the nutation phenomenon occurs (see supplementary video 1)

### Origin of *K*_0_ and regulation processes

The maintenance of hook curvature is achieved by establishing a spontaneous curvature (see Fig. 5B), in particular during the first stages of development. The plant can play on its spontaneous curvature via tropisms and differential elongation (8–10).

However the change of shape of the *Averrhoa carambola* leaf could not be explained by a switch from negative to positive gravitropism only. Chelakkot and Mahadevan (10) proposed a model where the interplay between weight, mechanics and growth zone creates this curvature. Here, we describe a different situation where spontaneous curvature–not elasticity–is important, especially in the beginning.

Gravitropism is a natural candidate to explain the hook shape. However, plants growing with impaired gravisensing still exhibit a hook. In our experiment with a clinostat, the hook is still present although the curvature is reduced by a factor 2 (Fig. 6). The very same effect was also observed in the case of the hypocotyl of *Pisum sativum* (22). Gravity would then be a clue rather than a cue (1). Miyamoto et al. (22) argue that the hypocotyl hook shape is already present in the grain. Leaves are however different: they can first grow unconstrained, quite straight, or even curved in the other direction. Still, in all cases, the plant is able to build and maintain a spontaneous curvature independently of any gravitropism. This first curvature can only be based on the adaxial-abaxial asymmetry of the leaf (23).

Remains the open question of what is regulated. For vine tendrils, where gravity can be neglected, (24) assumed that the spontaneous curvature relaxes to the actual curvature so that no elastic strain remains. In their model, Chelakkot and Mahadevan (10) hypothesize that spontaneous curvature is adjusted according to the real curvature. This idea was an extension of the autotropism idea initiated by Bastien et al. (8). Here we find that the spontaneous curvature in the growing zone is actively regulated (Fig. 6), and leads to a roughly constant real curvature profile (Fig. 3). We also observe the vanishing of curvature at the end of the growing zone. Outside of the growing zone, the process of lignification helps maintain a flat state. This behavior is consistent both with Goldstein and Goriely‘s hypothesis and Chelakkot and Mahadevan‘s framework. However for a large leaf with the presence of gravity, there is necessarily a gap between the actual curvature and the spontaneous one. In this way *K*_0_ can never be equal to *K*_*//*_. What could be regulated is the actual curvature *K*_*//*_, but the plant needs a way to perceive it. The remarkable point here is the time dependent evolution of this structure is controlled by the distance to the apex. Plants are sensitive to the total strain and not to the elastic stress (25). Therefore, the growing region which has the lowest Young’s modulus generates the highest elastic strains and is more prone to regulation. This is true if the weight moment is not too small i.e. we are not too close to the apex. The second important point is that elastic strain is proportional to *K*_*//*_ − *K*_0_ and not to *K*_*//*_ only. It is tempting to think that it is this quantity, rather than *K*_*//*_, that is perceived and regulated. Indeed, our results show that at later times displays a steady profile. Interestingly enough, the maximum of *K*_*//*_ − *K*_0_ consistently coincides with the zone of unbending (i.e. decreasing *K*_*//*_ values), at a fixed distance from the apex. This suggests that the strain due to self-weight could participate in triggering the straightening process. We can also look at the result of leaves grown on the clinostat, simulating a null gravity. We obtain that *K*_*//,clino*_ is reduced compared to *K*_*//*_ (see Fig. 6). Assuming similar flexural rigidity *B*, corresponding spontaneous curvatures can be deduced, and *K*_*//,clino*_ is also found to be around half *K*_0_ observed with gravity. It thus shows that there is a spontaneous curvature *K*_0_, even without gravity, and that with gravity, the spontaneous curvature, is larger. Based on our results, we propose that, at the beginning, the regulation is on the spontaneous curvature *K*_0_, since there is not effect of gravity yet, and at later times, when and if the gravity effect kicks in, the regulation bears on the perceived deformation *K*_*//*_ − *K*_0_. It is very likely that improving Chelakkot and Mahadevan‘s theoretical framework by doing the regulation on *K*_*//*_ − *K*_0_ would also not change the results qualitatively.

### Hook shape and hypocotyls

Compound leaves and hypocotyls are radically different organs. In the first place, their physiological functions completely differ. On anatomical grounds, hypocotyls also have an actual cylindrical symmetry when the rachis of compound leaves have a bilateral symmetry (see cross sections in Fig. S5). Their developmental contexts also differ. Hypocotyls first grow underground, in complete darkness, and light triggers the opening of their hook (26). Meanwhile the leaves of *Averrhoa carambola* leaves grow exposed to light, with no bud around them. Despite these differences, our study shows that *Averrhoa carambola* compound leaves not only share a hook shape with most dicotyledonous hypocotyls, but also their curvature and growth kinematics (2, 21). This profound resemblance may be a useful guide to investigate and understand leaves’ hook shape regulation at the microscopic scale. For example, leaf and hypocotyl hooks could be based on similar organizations of auxin efflux carriers or auxin and ethylene distribution, two important actors in the regulation of hypocotyl hooks (4).

Finally, let us mention the apical hook of hypocotyls has been shown to increase the survival rate of rice seedlings (27) as it might help to protect the cotyledons which are the initial source of nutrients of the frail seedling. In the case of compound leaves, the context remains different and the question of the function of the apical hook remains open.

## Conclusion

Associating quantitative kinematics observation with a simple mechanical test, we demonstrated in this article that the downward curve shape of developing compound leaf shape is an active and regulated process. By reversing the arrow of gravity we show that rigidity and spontaneous curvature are sufficient to compensate gravity effects, especially at the beginning of development. We find that this process is regulated in space and time, first with a fixed spontaneous curvature at the tip, when the deformation due to gravity of the small leaf is quasi-inexistent. When gravity starts to be important with the larger size of the leaf, we then observe fixed deformation at a given distance from the tip, meaning that the spontaneous curvatures is then regulated to achieve this deformation. In this way the leaf uses the gravity effect to tune its own regulation. Our findings bring experimental support to the recently developed theoretical framework mixing autotropism and mechanics which hypothesized a regulation of natural curvature (10). We have additionally shown that gravitropism has a limited influence in the processes involved in the regulation of the leaf’s shape. These findings in the compound leaves’ hook are analogous to what has been observed on the hypocotyls’. As in stem growth, we found the localization of the growth zone situated at a fixed distance from the apex, as well as the change in the mechanical properties of the tissues along the rachis during development. When the rachis becomes rigid enough not to be deformed by gravity anymore, it reaches locally its final straight shape. We can hypothesise that the curving of the rachis is to probe the gravity effect, and finally use it in order to converge toward the right straight shape (10, 28).

## ACKNOWLEDGEMENTS

The clinostat used for these experiments has been designed and built by Mathieu Receveur. We would like to thank Amina Saadani, Lea Laffont and Clara Billard for their help with preliminary experiments. We are grateful to Bruno Moulia for inspiring the “flipping test” idea. We thank Richard S. Smith and Olivier Pouliquen for their insightful suggestions and comments. We would like to thank Emmanuel de Langre and Olivier Hamant for continuous feedback on our research, and also Cyprien Gay for helpful discussions about mechanics. MR is thankful to “Ecole Doctorale Frontières du Vivant (FdV) Programme Bettencourt” for financial support. The authors declare no conflict of interests.

## Footnotes

1 a rotating device usually dedicated to disrupt gravisensing in plants.

2 *L* actually shows some degree of oscillation in Fig. 3. This corresponds to curvature oscillations orthogonal to the principal plane usually referred as nutation (1). Here we disregard nutation as its amplitude is negligible compared to the unfurling motion. However, the emergence of the final shape is a continuous and 3D phenomena. Splitting this process into several motions is convenient but remains a convention. We will address nutation in a future study.

