## Supplemental figures for "Hook shape of growing leaves results from an active regulation"

### SUPPLEMENTARY DATA

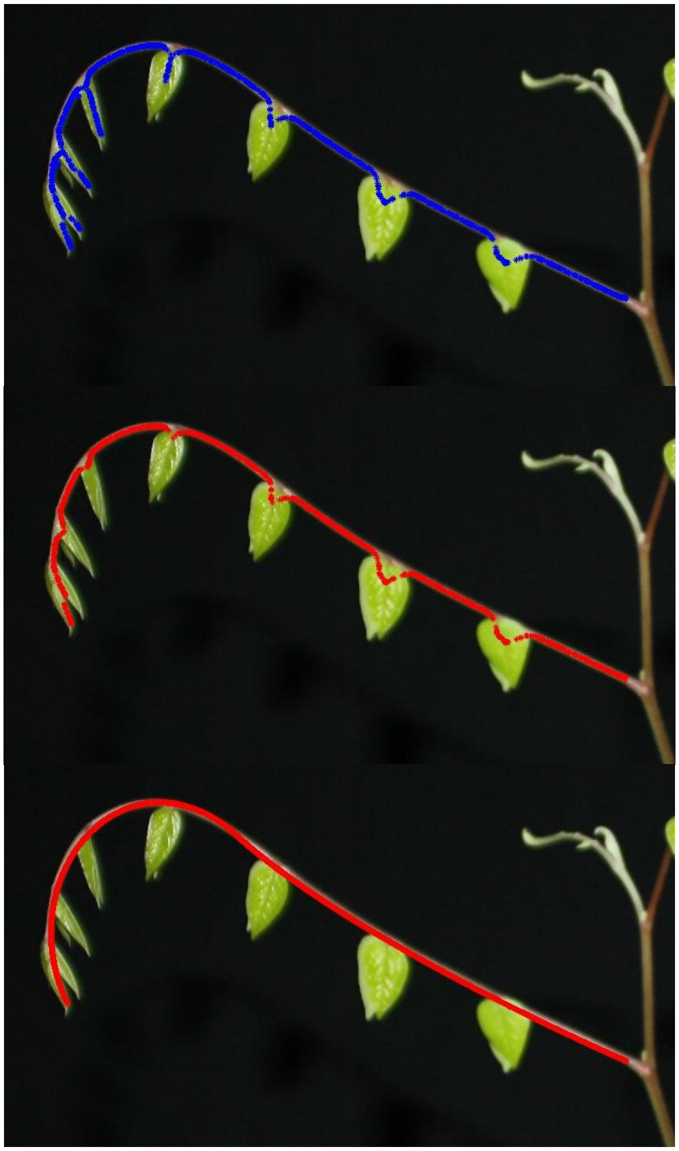

**Fig. S1:** Skeletonization procedure for growth kinematics acquisitions. (a) Blue crosses correspond to the output of the Voronoi-based algorithm. The retrieved skeleton does not perfectly correspond to the petiole of the leaf. It is highly affected by the presence of the leaflets. (b) Main branch of the previously obtained skeleton. Here, this first step allowed to get rid of most deformations due to the apical leaflets. (c) Finally, the main branch of the skeleton is smoothed by fitting it with Bézier curves. The final result matches the rachis of the leaf in a much more satisfying way.

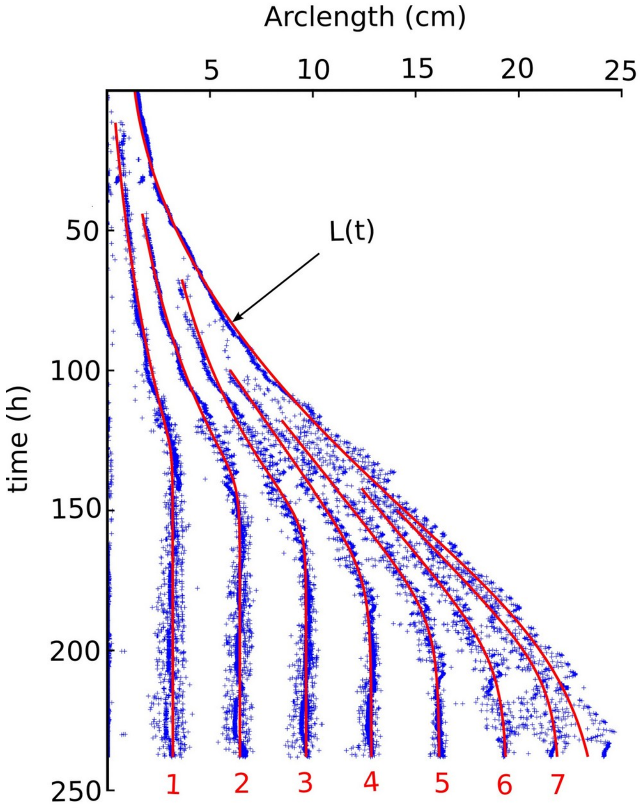


**Fig. S2:** Leaflets trajectories smoothed by analytical fit inspired by typical logistic growth laws. We found that the raw trajectories (blue crosses) were nicely fitted by a sigmoid function (red lines)

For leaflets 1 to 3:

$$s_f(t)=\left[\left(s_1+s_2\cdot e^{t/\tau}\right)^{-\gamma}+s_{sat}^{-\gamma}\right]^{1/\gamma}$$

For leaflets 4 to 7:

$$s_f(t)=\left[\left(s_1+s_2\cdot t\right)^{-\gamma}+s_{sat}^{-\gamma}\right]^{1/\gamma}$$

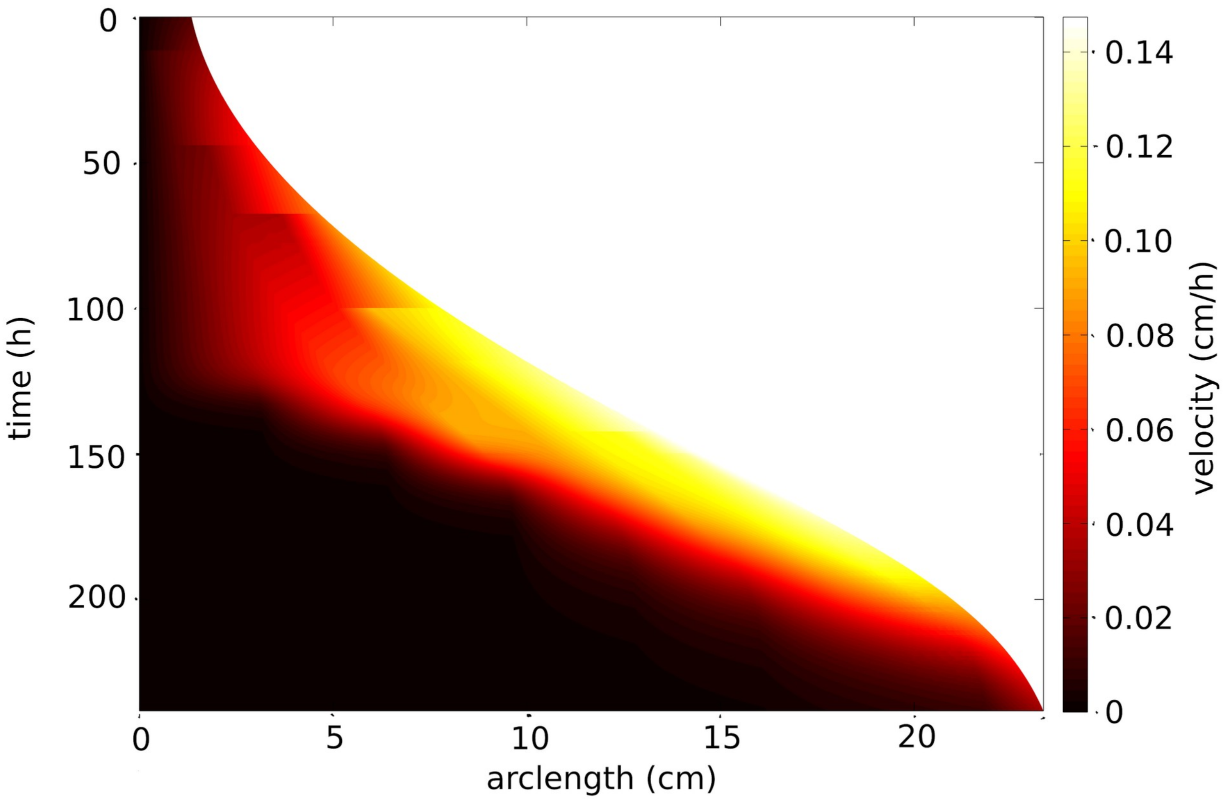


**Fig. S3**: Interpolation of the growing zone. An effective velocity of leaflets has been interpolated between leaflets. Note the colocalization of this zone with the bent zone evidenced in Fig. 3.

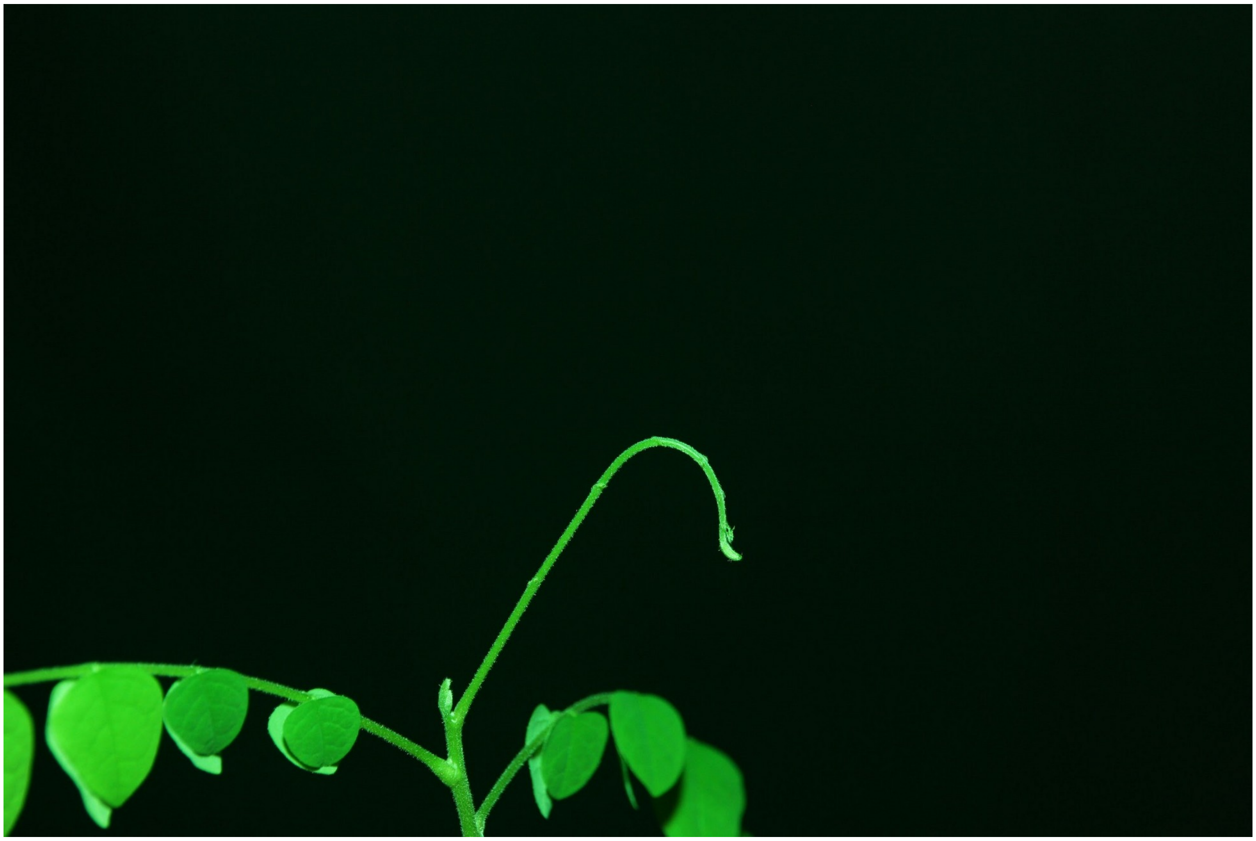

**Fig. S4**: Example of an *Averrhoa carambola* leaf growing without its leaflets. The leaflets of this leaf have been cut off with tweezers, except for the most apical ones that are too delicate to withdraw at first. We see that the typical hook shape is still present. The unfurling motion is also preserved and the whole development of the leaf is qualitatively comparable to the one of a normal leaf.

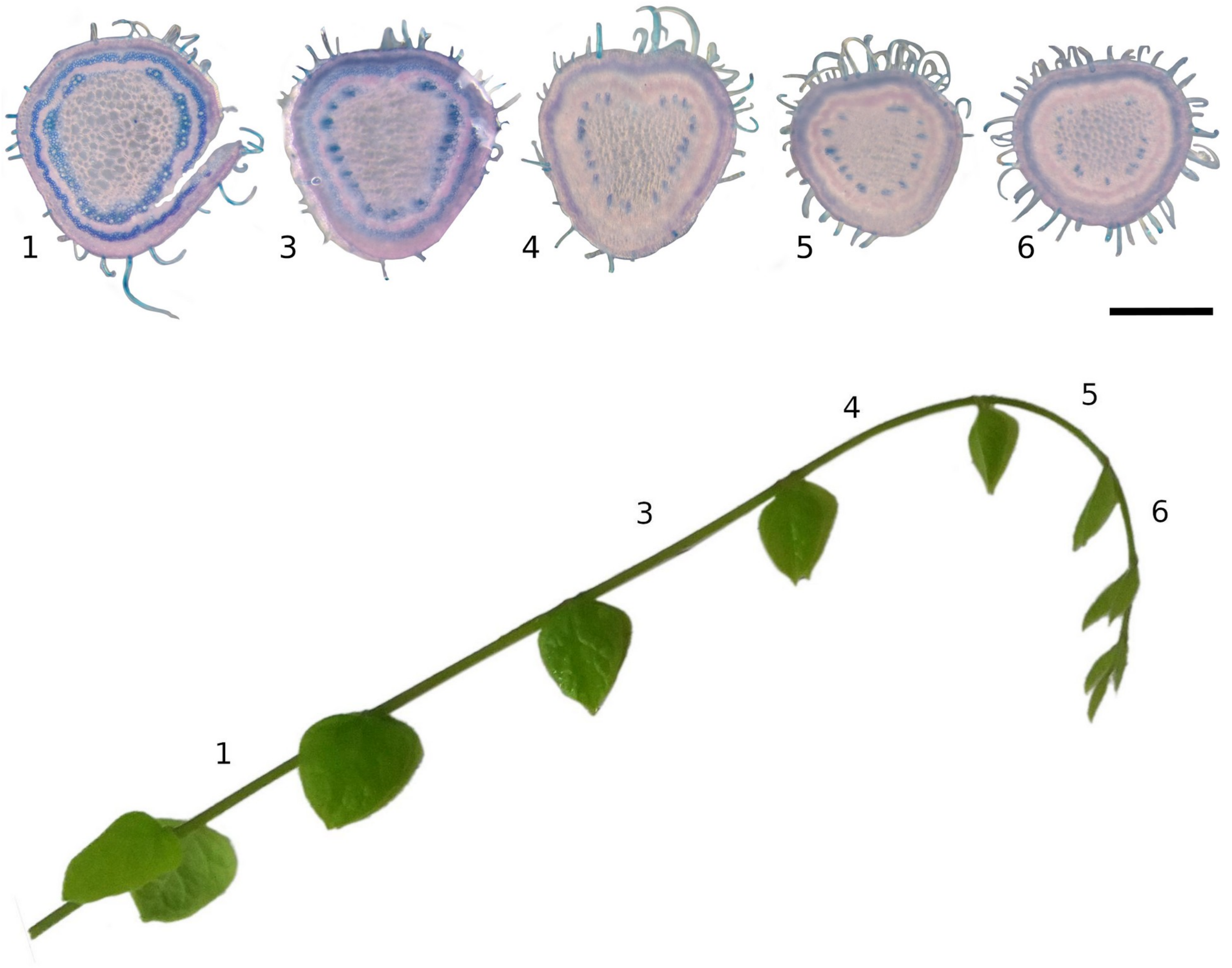


**Fig. S5:** Lignification status of the rachis. (top) Cross-sections of the rachis colored with the dual carmine-iodine stain. Blue zones indicate lignified tissues, pink-red zones indicate non-lignified tissues. Samples are ordered from the base (left) to the apex (right). Scalebar is 500 micrometres. (bottom) A quantification of the lignification rate for each cross-section (ratio of lignified area over total area). The score is color coded on each interleaflet.

***Gradient of lignification from apex to base***

The simple carmine and iodine green dual staining reveals --- qualitatively --- an obvious gradient of lignification along the rachis (see supplemental Fig. 5). Near the apex, the lignification index is 7% and progressively increases up to 17% at near the base. Observing the successive samples shown, from apex to base, we trace the path of the lignification process. At the apex, we see the presages of a ring of bast fibers. Its development starts on the adaxial face of the rachis and then progressively propagates to the abaxial one. These bast fibers eventually form a complete ring. We also see that, near the apex, some vessels are already lignified. The number of these vessels increase as we go towards the base. It looks like, unlike for bast fibers, vessels develop from abaxial face to the adaxial one. Finally, from these vessels, we see that a ring of xylem develops.
